# Tumor IsomiR Encyclopedia (TIE): a pan-cancer database of miRNA isoforms

**DOI:** 10.1101/2020.08.20.259713

**Authors:** Xavier Bofill-De Ros, Brian Luke, Robert Guthridge, Uma Mudunuri, Michael Loss, Shuo Gu

## Abstract

MicroRNAs (miRNAs) function as master regulators of gene expression in many physiological and pathological conditions including cancer. Sequence variants or isoforms (isomiRs) can account for between 40 to 60% of total miRNA counts, yet despite this overwhelming abundance, their function continues to be debated. Recent studies demonstrate that certain isomiRs can regulate unique sets of target mRNAs by altering their seed sequence or stabilizing 3’ pairing, while others are decay intermediates indicating an active miRNA turnover. Given their short sequence length and high heterogeneity, mapping isomiRs can be challenging; without adequate depth and data aggregation, low frequency events are often disregarded. To address these challenges, we present the **<u>T</u>**umor **<u>I</u>**somiR **<u>E</u>**ncyclopedia (TIE): a dynamic database of isomiRs from over 10,000 adult and pediatric tumor samples in The Cancer Genome Atlas (TCGA) and The Therapeutically Applicable Research to Generate Effective Treatments (TARGET) projects. A key novelty of TIE is its ability to annotate heterogeneous isomiR sequences and aggregate the variants obtained across all samples and datasets. The database provides annotation of templated and non-templated nucleotides as well as other advanced analysis. All data can be browsed online or downloaded as simple spreadsheets. Here we show analysis of isomiRs of miR-21 and miR-30a to demonstrate the utility of TIE. TIE search engine and data are hosted at https://isomir.ccr.cancer.gov/.

## Introduction

MicroRNAs (miRNAs) are a class of non-coding RNAs (~23nt) with important regulatory functions [1]. Since their discovery two decades ago, thousands of miRNA genes have been identified in viral, plant and animal genomes [2]. By binding to the Argonaute protein, miRNAs form the RNA-induced silencing complex (RISC) which modulates the expression of the vast majority of human transcripts [3,4]. Dysregulation of miRNAs has been linked to many diseases including cancer [5,6].

Each mature miRNA can present as a variety of isoforms (isomiRs) that differ in length and sequence composition [7,8]. When isomiRs gain or lose nucleotides at the 5’ end, they are called 5’ isomiRs; those which vary at the 3’ end are 3’ isomiRs. We refer to 3’ isomiRs that lose nucleotides as “trimmed” and those that gain non-templated nucleotides as “tailed”. There are multiple origins of isomiRs which reflect dynamic events that occur during biogenesis and post-maturation. During biogenesis, the cleavage sites of DROSHA and DICER1 define the ends of mature miRNAs [9]. Cleavage by DROSHA or DICER1 at alternative sites generates “templated” isomiRs - sequences that match the genomic reference but vary in start and/or end positions [10–12]. Furthermore, isomiRs can arise from post-maturation 3’ editing - ribonucleases remove nucleotides from the 3’ end whereas terminal nucleotidyl transferases (TENTs) incorporate non-templated nucleotides, resulting in trimmed or tailed isomiRs respectively [13]. Unlike the 5’ end of a miRNA which tightly binds to Argonaute, the 3’ end is often accessible, making 3’ modification a frequent event [14,15]. As a result, 3’ isomiR is the most prevalent type, both in terms of the number of miRNAs displaying these variations and overall abundance.

Multiple lines of evidence suggest that isomiRs possess unique biological function: isomiRs are evolutionarily conserved and their expression is tightly regulated. Case studies indicate that naturally existing isoforms can have distinct activities across a wide range of biological processes, including regulation of cytokine expression, facilitation of virus proliferation, and promotion of apoptosis [13]. Mechanistically, 5’ isomiRs present a different seed composition than the corresponding canonical miRNAs and therefore regulate an alternative set of mRNAs. Sequence modifications at the 3’ end do not change the seed sequence, but nonetheless are reported to play critical roles in regulating miRNA targeting specificity and stability [15–18].

It is well-documented that miRNAs plays a critical role in cancer development [19]. Expression profiles of miRNAs, together with other transcriptomic and genomic approaches, have contributed to the characterization of the main changes and commonalities underlying different tumors and tumor subtypes [20,21]. Interestingly, it has been shown that alteration in isomiR profiles (rather than overall miRNA abundance) correlates with cancer progression, which suggests a unique role for isomiRs in tumorigenesis [22,23]. Multiple reports suggest that isomiRs can be used as biomarkers for tumor detection, classification and patients prognosis [24–27]. These observations highlight the importance of having computational tools to analyze isomiR expression profiles from sequencing data to further investigate their functions.

However, it is difficult to investigate isomiRs with existing databases. While miRBase [28], MirGeneDB [29] and other databases list evolutionarily conserved isoforms across different species (including human, mouse, fly and others) [30,31], comprehensive documentation of isomiRs in cancer samples remains missing. The main challenge in the study of isomiRs is the extreme heterogeneity in their abundance and sequence composition. Various combinations of 5’ and 3’, templated and non-templated, and trimmed and tailed isomiRs exist together. While the overall 3’ isomiRs could be dominant (>50%) for certain miRNAs, the abundance of a large portion of the individual isomiRs is close to the level of sequencing noise. As a result, many low frequency isomiRs are often discarded and remain undetectable with prevalent mapping algorithms. This highlights the need for specialized algorithms capable of mapping and efficiently processing all isomiRs in order to comprehensively analyze cancer isomiRs [32,33].

Here, we aim to provide the field with a database that contains a complete list of isomiRs from the largest existing cohort of human tumor samples. Towards this, the TIE database analyzes ~97 billion reads and ~16 billion small RNA sequences provided by The Cancer Genome Atlas (TCGA) and The Therapeutically Applicable Research to Generate Effective Treatments (TARGET) projects, and reports isomiRs in a high-throughput and automated manner (Figure 1A). The depth and diversity of the TIE database allow the comprehensive study of highly specific modifications. Furthermore, we provide easy download access to all files and reports generated to allow users with specific questions to focus on unique aspects of the isomiR biology. TIE search engine and database are hosted at https://isomir.ccr.cancer.gov/.

**Figure 1.**
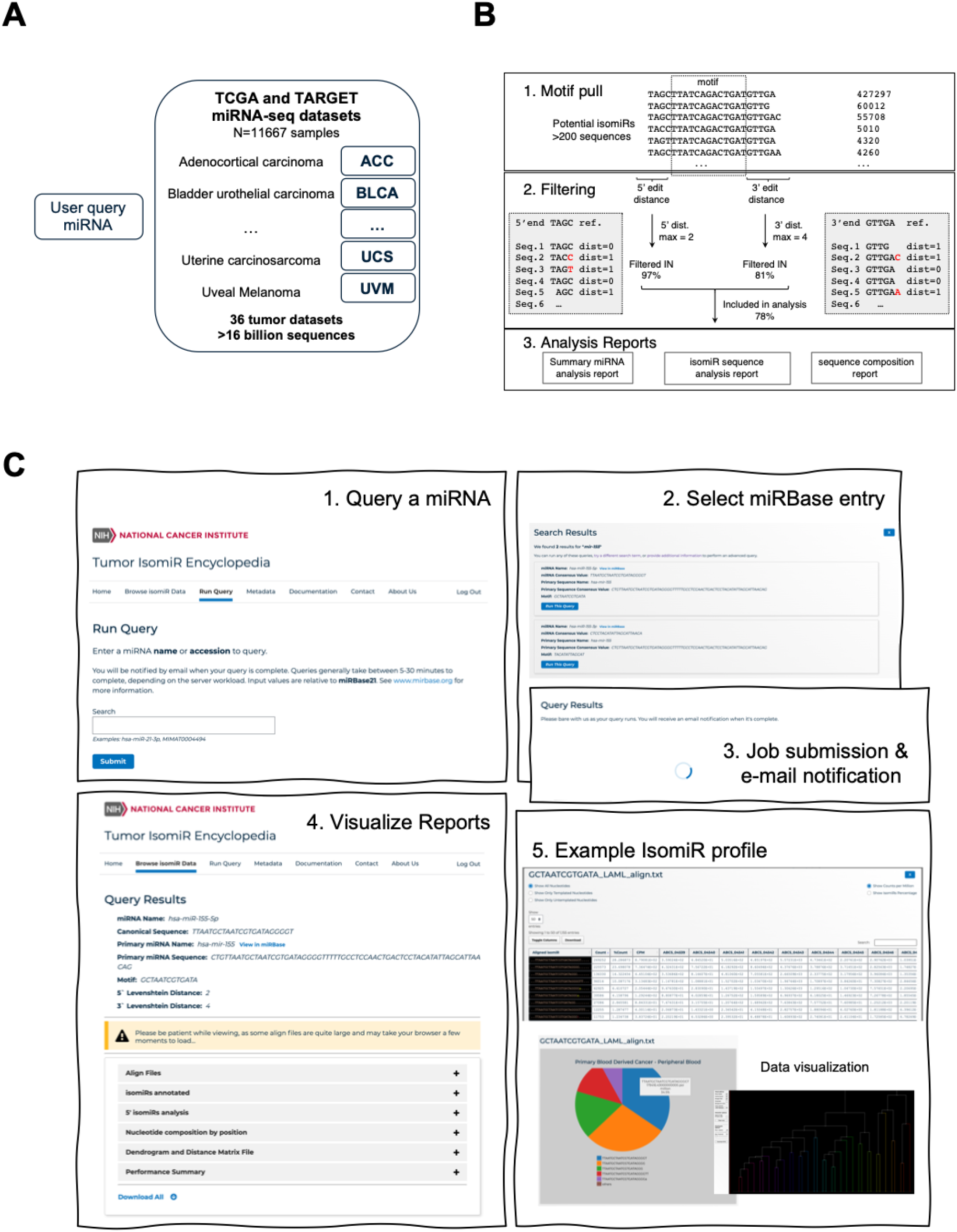
Structure of the Tumor IsomiR Encyclopedia algorithm and outputs. (***A***) Schematic of the data structure and miRNA-seq datasets analyzed. (**B**) Algorithm used in TIE analysis and example of the mapping flow. (**C**) Features of TIE database and basic steps in isomiR analysis.

## Method

### Mapping algorithm and analysis of NGS data

TIE uses the QuagmiR algorithm, previously published by our group[34], to align miRNAs and their isoforms (Figure 1B). Briefly, this alignment method facilitates mapping of heterogeneous isomiRs to the reference miRNA by allowing a different number of mismatches at the 5’ and 3’ ends to reflect differences in the nature of their biogenesis. This algorithm has two steps: 1) Reads are pulled based on a unique motif (13-mer) that distinguishes between the miRNA of interest and other miRNAs from the same family or others with high sequence similarity (eg. miR-148a and miR-148b). The current set of 13-mer motifs is able to distinguish close to 95% of miRNA sequences [34]. 2) Edit distances (Levenshtein) between the canonical miRNA sequence and each pulled read are calculated for the segments upstream (5’ edit distance) and downstream (3’ edit distance) of the motif. TIE uses by default a maximum 5’ edit distance of two and 3’ edit distance of four. Further documentation on the TIE database and its underlying algorithm can be found at the GitHub wiki (https://github.com/Gu-Lab-RBL-NCI/TIE-database/wiki).

NGS data from TCGA and TARGET projects were downloaded as bam files, converted to fastq format and then summarized into collapsed files, which were stored on the server and used as input for an isomiR query.

## Results

### Data content and usage

The TIE database dynamically aligns the small-RNA-seq data from TCGA and TARGET. These data were collected from 11,667 patient samples across 36 types of cancer (33 adult tumors and 3 pediatric tumors). Metadata of these samples can be found in the corresponding tab on the TIE site. A brief summary is listed in Supplementary Table 1. TIE enables users to query the isomiR profile of any miRNA documented in the miRBase database. For each miRNA, isomiR profiles of cancer samples are compared to those from corresponding normal tissues and shown as pie-charts. Detailed isomiR analyses for each patient sample are reported as column-wise tables that are easily downloaded as spreadsheets. Once files have been downloaded, users can perform additional analysis with Microsoft Excel, Python, R or Perl, among others.

Contents of the TIE database can be searched and downloaded with a web browser. Users can either browse previously analyzed isomiR profiles or perform a new query. The TIE database output includes reports on isomiR expression and sequence composition among other parameters. A screenshot of the website is found in Figure 1C. Below we describe the use of the website and details of the outputs, and provide sample analyses of miR-21 and miR-30a.

### Browse isomiR data

IsomiR profiles of many highly expressed miRNAs were precompiled and ready to download. This page serves as the entry point for users to search and browse reports stored on the online servers. For each report, users are provided with a brief description including the miRNA name, mature sequence, corresponding pri-miRNA sequence and the unique motif used in the alignment.

### Run query

This page guides the users to run a *de novo* isomiR analysis for their miRNA of interest across all cancer types. Once the task is completed, the results will be displayed on the same url page (Figure 1C). If the users provide their email, they will be notified once the task is complete.

### Align files

A primary aim of the TIE database is to aggregate all miRNA isoforms detected in human tumors. The “Align files” page is an entry point for browsing the sequence variability of any given miRNA across this big pan-cancer dataset. This page displays the sequence and abundance of all isomiRs for the querying miRNA across all cancer types. Templated nucleotides, which can be mapped back to the genome, are displayed in uppercase text and non-templated nucleotides are shown in lowercase with a unique color designated for each non-templated nucleotide. The dot notation is used as a position holder for alignment to the pri-miRNA sequence. Users can filter the results to visualize only isomiRs with templated nucleotides or non-templated nucleotides. The absolute amount and relative percentage of each isomiR are reported for each patient sample and for each cancer type.

### IsomiRs annotated

To facilitate the study of isomiRs in tumors, the “isomiRs annotated” page tabulates detailed annotation and analysis of each isomiR. Some of the features reported include read length, number of nucleotides trimmed, number of nucleotides tailed, variations at the 5’ end, tail sequence, seed sequence and difference from the canonical miRNA measured as edit distances. In addition, abundance is reported in absolute counts (CPM) and relative percentages. Also, TIE provides the option to compare with dynamic pie charts the main isomiRs between different cancer types or between the different sample types (Normal, Tumoral and Metastasis). This allows the users to perform more specific analysis of the downloaded data on specific groups such as uridylated, adenylated or trimmed isomiRs.

### 5’ isomiRs analysis

The function of a miRNA is largely defined by its seed sequence, which is a stretch of nucleotides from positions 2 to 7. Recent literature has shown that cleavage by either DROSHA or DICER1 at alternative sites can generate 5’ isomiRs with altered seed sequences [11]. This process is known as seed shift. The mechanisms regulating seed shift are not yet fully understood, but include aspects such as changes in the secondary structure of miRNA precursors [10–12] and/or the recruitment of RNA-binding proteins[35]. The “5’ isomiRs analysis” section facilitates the analysis of this phenomenon in the pan-cancer datasets. Users can find reports of the 5’ Fidelity index and the Effective Number of seeds. The 5’ Fidelity index is an average of the cleavage site position offset weighted by the number of supporting reads, while the Effective Number of seeds is calculated as an inverted Simpson index. More details of their calculation and interpretation can be found in the online documentation of the database.

### Nucleotide composition by position

It has been determined that certain internal RNA editing events occur in miRNAs. For instance, ADAR enzymes catalyze the A-to-I conversion [36], thus altering miRNA targeting in tumors [37,38]. In NGS results, inosine is read as the misincorporation of a G, while m^1^A methylation is read as the misincorporation of random nucleotides with a concurrent drop in the sequence coverage [39]. In order to identify such editing events, as well as other templated and non-templated incorporations on the 5’ and 3’ ends, the section “Nucleotide composition by position” reports the percentage of each nucleotide found in each position in all the aligned isomiRs.

### Dendrogram and Distance Matrix Files

This section uses the previously reported data to group and classify tumoral samples and cancer types according to the distribution of their miRNA isoforms (isomiR profiles). Complete distance matrices for each dataset are provided, as well as a graphical visualization of the different cancer types in a dendrogram.

### Performance Summary

This section provides technical information about the performance of the motif search used for isomiR detection across the different datasets. It informs the user on the adequacy of the settings used for their specific query. Some of the parameters reported are the number of samples in which the queried isomiR (motif) can be found, minimum and maximum variation (measured as edit distances) between isomiRs and the canonical miRNA, as well as sequence diversity within matched isomiRs.

### Case study: isomiR analysis in a pan-cancer datasets

One of the most well studied miRNAs with oncogenic function is miR-21-5p [40]. Analysis of its expression across different tumors and corresponding normal samples collected in the TIE database confirms previous reports of its general overexpression in cancer (Figure 2A). miR-21 is upregulated ~7 fold in the kidney papillary cell carcinoma (KIRP), while a modest increase of miR-21 (~2 fold) was observed in other cancers including lung squamous cell carcinoma (LUSC) and prostate adenocarcinoma (PRAD). Despite the well-established upregulation of miR-21 in tumors, it remains unclear to what extent the isomiR profile changes during tumorigenesis; study of this will lead to not only novel biomarkers, but also to the identification of potential regulations of miR-21 biogenesis and decay. To this end, we analyzed miR-21-5p isomiRs with TIE (Figure 2B). Although the average percentage of isomiRs oscillates between 24% and 38% in different cancers, miR-21 isomiR is more abundant than the canonical miR-21 in many patient samples. Interestingly, the level of isomiRs is significantly higher in some cancer types such as lung squamous cell carcinoma (LUSC) when compared to their corresponding normal samples (LUSC-N). Further analysis revealed that the mono-adenylated isoform of miR-21 was enriched in tumor samples (Figure 2B). These results support a hypothesis that certain miR-21 isomiRs, in particular the mono-adenylated isoform, may play a functional role during tumorigenesis.

**Figure 2.**
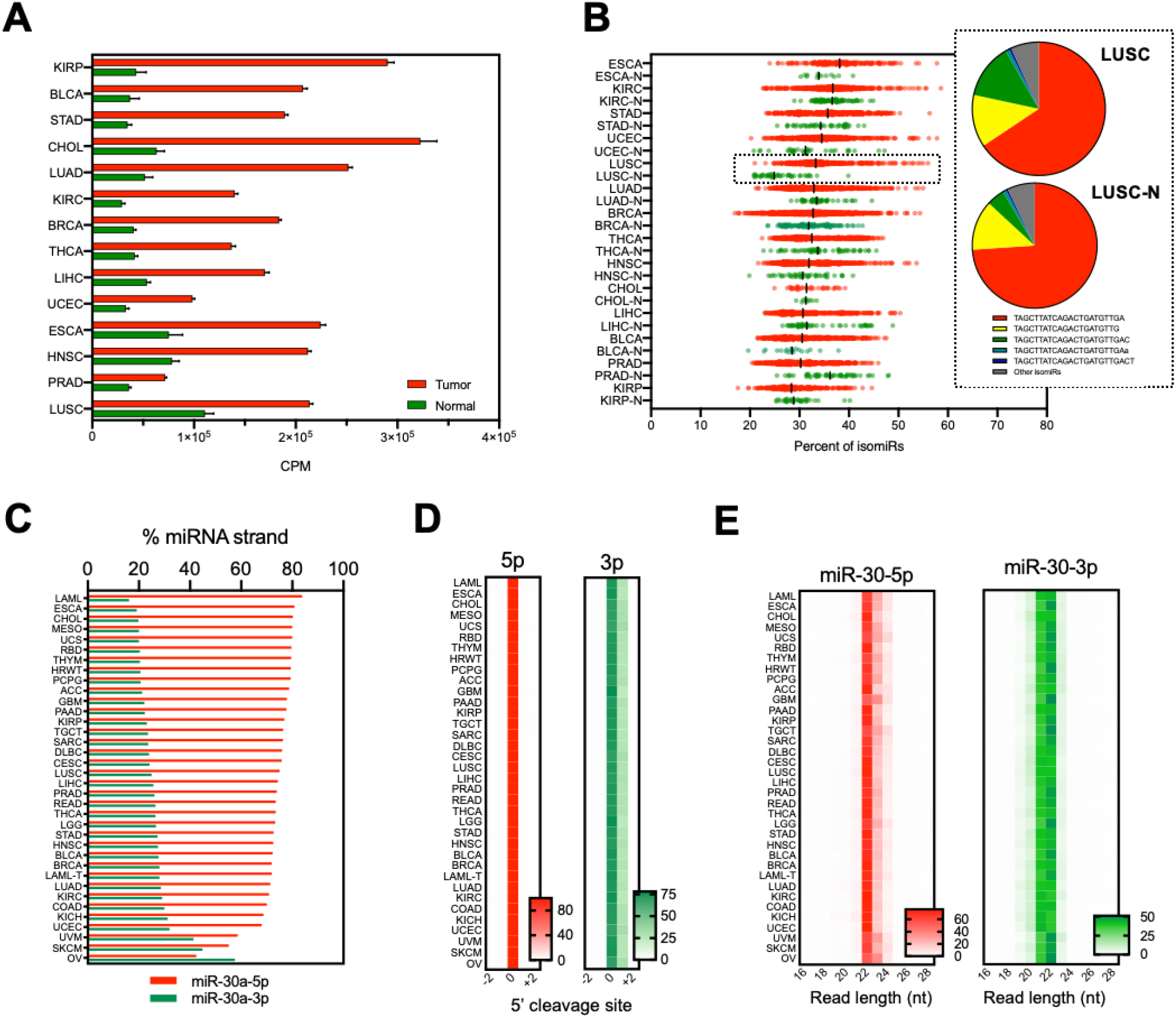
Case study of isomiR analysis in a pan-cancer dataset. (**A**) Expression of miR-21-5p (CPM) in different cancer types and corresponding normal tissues. (**B**) Scatter plot of the percentage of non-canonical isomiRs of miR-21-5p in each sample within different cancer types and corresponding normal tissues (N). The black line indicates the median of the sample distribution. The snippet shows an example of the relative abundances of the canonical miR-21-5p (in red) and other dominant isoforms in lung squamous cell carcinoma (LUSC) and corresponding normal solid tissue (LUSC-N). (**C**) Plot of the average relative 5p and 3p strand abundance for miR-30a in different cancer types. (**D**) Heatmap with the 5’ cleavage site of miR-30a-5p and miR-30a-3p. The canonical cleavage site is indicated as 0; other sites are indicated by the number of nucleotides downstream (−) or upstream (+). Abundance of isoforms starting at each position is indicted as a relative percentage. (**E**) Heatmap with the length distribution of all isoforms of miR-30a-5p and miR-30a-3p (relative percentage).

Another interesting example that highlights the complex regulation of miRNA biogenesis is arm switching, during which the functional strand joining Argonaute (guide strand) and the strands being discarded (passenger strand) are switched. This mechanism is regarded as a major driver of miRNA evolution [41] and one of the signatures of miRNA dysfunction in tumors [42,43]. Despite many specialized tools for analyzing miRNA arm switching[44], pan-cancer analysis of this phenomenon was not available. TIE offers a unique opportunity to study arm switching in tumors and to investigate the possible role of isomiRs in this process. One of the most well-known examples of arm switching occurs in miR-30 [42]. We therefore analyzed the relative abundances of miR-30a 5p and 3p strands in each tumor type (Figure 2C). While the 5p strand is in general the dominant strand, a remarkable increase in the levels of 3p over 5p (arm switching) was observed in uveal melanoma, skin cutaneous melanoma and ovarian serous adenocarcinoma (UVM, SKM and OV), suggesting a functional role for arm switching in these cancers. Interestingly, while DROSHA and DICER1 cut at the same positions during biogenesis (Figure 2D), longer isomiRs of the miR-30 5p and 3p strand were enriched in UVM, SKM and OV (Figure 2E). This suggests that post-maturation modifications (such as 3’ tailing) could be the underlying mechanism of the observed arm switching [45].

## Discussion

It is clear that miRNAs play essential roles in cancer development, and that miRNA function is largely determined by sequence and abundance. IsomiRs are highly prevalent as a result of sequence modifications, however despite decades of research into the functions of miRNAs in cancer, little is known about isomiR biogenesis and function in tumors. Recent studies have shown that isomiRs have distinct targeting specificities and/or turnover rates [15–18], suggesting that they may play important roles during tumorigenesis. In fact, it was shown that the oncomiR miR-21 is regulated via TENT4B adenylation and subsequent degradation by the exonuclease PARN[46], a mechanism implicated in the regulation of p53 levels [47]. In addition, TUT4/7 uridylation of miR-324 induces DICER1 miscleavage and strand switching, and results in the production of an aberrant miR-324 isomiR, which in turn promotes cell proliferation in glioblastoma [35]. Similarly, tailing and trimming accompanies strand switching in miR-574 and contributes to gastric cancer progression. The miR-200 family is a key regulator of the epithelial to mesenchymal transition by targeting ZEB1 [48]. Recent analysis shows that high levels of ADAR1 editing in the seed region of miR-200b in thyroid carcinomas play an important role in proliferation, invasion, and migration [49,50]. These examples support the idea that isomiRs are involved in the generation of novel oncogenic isomiRs or the loss of function of well-established tumor suppressor miRNAs.

Similar to other databases [28–31], TIE accurately reports the expression of miRNA isoforms. However, TIE excels at providing detailed characterization and extensive analyses of 5’ and 3’ isomiRs. It also provides easy access to the most comprehensive set of adult and childhood cancers. This enables researchers to perform big data analysis without downloading or storing sensitive data locally. The big data analysis of isomiRs in pan-cancer datasets could pave the way for a better understanding of their role in tumorigenesis. TIE offers an excellent opportunity for researchers working on isomiR functions to characterize different isomiR groups and their cancer of interest.

One limitation of the current version of TIE is that the settings regarding the motif and the number of mismatches tolerated are pre-determined. As a result, the isomiR analysis is restricted to the well annotated miRNAs in the miRBase. However, recent studies have identified hundreds of novel miRNAs which might be only expressed in specific cells [51]. Future versions of TIE will allow advanced users to take advantage of a flexible search algorithm and query the isomiR profiles of unannotated or viral miRNAs. It will be also interesting to explore the impact of isomiRs on cancer by combining TIE with other datasets such as the exome sequencing or the transcriptomic profiles available for both TCGA and TARGET.

## Supplementary Tables

**Supplementary Table 1:**

| Tumor Description | Dataset | N | Sample types |
| --- | --- | --- | --- |
| ADRENOCORTICAL CARCINOMA | ACC | 80 | Primary Tumor |
| BLADDER UROTHELIAL CARCINOMA | BLCA | 437 | Primary Tumor, Solid Tissue Normal, Metastatic |
| BREAST INVASIVE CARCINOMA | BRCA | 1207 | Primary Tumor, Solid Tissue Normal, Metastatic |
| CERVICAL SQUAMOUS CELL CARCINOMA AND ENDOCERVICAL ADENOCARCINOMA | CESC | 312 | Primary Tumor, Solid Tissue Normal, Metastatic |
| CHOLANGIOCARCINOMA | CHOL | 45 | Primary Tumor, Solid Tissue Normal |
| COLON ADENOCARCINOMA | COAD | 465 | Primary Tumor, Solid Tissue Normal, Metastatic, Recurrent Tumor |
| LYMPHOID NEOPLASM DIFFUSE LARGE B-CELL LYMPHOMA | DLBC | 47 | Primary Tumor |
| ESOPHAGEAL CARCINOMA | ESCA | 200 | Primary Tumor, Solid Tissue Normal, Metastatic |
| GLIOBLASTOMA MULTIFORME | GBM | 5 | Solid Tissue Normal |
| HEAD AND NECK SQUAMOUS CELL CARCINOMA | HNSC | 569 | Primary Tumor, Solid Tissue Normal, Metastatic |
| KIDNEY CHROMOPHOBE | KICH | 91 | Primary Tumor, Solid Tissue Normal |
| KIDNEY RENAL CLEAR CELL CARCINOMA | KIRC | 616 | Primary Tumor, Solid Tissue Normal, Additional New Primary |
| KIDNEY RENAL PAPILLARY CELL CARCINOMA | KIRP | 326 | Primary Tumor, Solid Tissue Normal, Additional New Primary |
| ACUTE MYELOID LEUKEMIA | LAML | 188 | Primary Blood Derived Cancer - Peripheral Blood |
| BRAIN LOWER GRADE GLIOMA | LGG | 530 | Primary Tumor, Recurrent Tumor |
| LIVER HEPATOCELLULAR CARCINOMA | LIHC | 425 | Primary Tumor, Solid Tissue Normal, Recurrent Tumor |
| LUNG ADENOCARCINOMA | LUAD | 567 | Primary Tumor, Solid Tissue Normal, Recurrent Tumor |
| LUNG SQUAMOUS CELL CARCINOMA | LUSC | 523 | Primary Tumor, Solid Tissue Normal |
| MESOTHELIOMA | MESO | 87 | Primary Tumor |
| OVARIAN SEROUS CYSTADENOCARCINOMA | OV | 499 | Primary Tumor, Recurrent Tumor |
| PANCREATIC ADENOCARCINOMA | PAAD | 183 | Primary Tumor, Solid Tissue Normal, Metastatic |
| PHEOCHROMOCYTOMA AND PARAGANGLIOMA | PCPG | 187 | Primary Tumor, Solid Tissue Normal, Metastatic, Additional New Primary |
| PROSTATE ADENOCARCINOMA | PRAD | 551 | Primary Tumor, Solid Tissue Normal, Metastatic |
| RECTUM ADENOCARCINOMA | READ | 165 | Primary Tumor, Solid Tissue Normal, Recurrent Tumor |
| SARCOMA | SARC | 263 | Primary Tumor, Recurrent Tumor, Metastatic |
| SKIN CUTANEOUS MELANOMA | SKCM | 452 | Primary Tumor, Solid Tissue Normal, Additional Metastatic |
| STOMACH ADENOCARCINOMA | STAD | 491 | Primary Tumor, Solid Tissue Normal |
| TESTICULAR GERM CELL TUMORS | TGCT | 156 | Primary Tumor, Additional New Primary |
| THYROID CARCINOMA | THCA | 573 | Primary Tumor, Solid Tissue Normal, Metastatic |
| THYMOMA | THYM | 126 | Primary Tumor, Solid Tissue Normal |
| UTERINE CORPUS ENDOMETRIAL CARCINOMA | UCEC | 579 | Primary Tumor, Solid Tissue Normal, Recurrent Tumor |
| UTERINE CARCINOSARCOMA | UCS | 57 | Primary Tumor |
| UVEAL MELANOMA | UVM | 80 | Primary Tumor |
| HIGH RISK WILMS TUMOR | HRWT_TARGET | 138 | Primary Tumor, Solid Tissue Normal, Metastatic, Recurrent Tumor |
| ACUTE MYELOID LEUKEMIA | LAML_TARGET | 369 | Primary/Recurrent Blood Derived Cancer (Peripheral Blood, Bone Marrow). Bone Marrow Post-treatment |
| RHABDOID TUMOR | RDB_TARGET | 78 | Primary Tumor, Solid Tissue Normal, Others |

## References

1 Gebert, L.F.R. and MacRae, I.J. (2019) Regulation of microRNA function in animals. Nat. Rev. Mol. Cell Biol. 20, 21–37

2 Kozomara, A. and Griffiths-Jones, S. (2014) miRBase: annotating high confidence microRNAs using deep sequencing data. Nucleic Acids Res. 42, D68–73

3 Friedman, R.C. et al. (2009) Most mammalian mRNAs are conserved targets of microRNAs. Genome Res. 19, 92–105

4 Eisen, T.J. et al. (2020) MicroRNAs Cause Accelerated Decay of Short-Tailed Target mRNAs. Mol. Cell 77, 775–785.e8

5 Sayed, D. and Abdellatif, M. (2011) MicroRNAs in development and disease. Physiol. Rev. 91, 827–887

6 Lin, S. and Gregory, R.I. (2015) MicroRNA biogenesis pathways in cancer. Nat. Rev. Cancer 15, 321–333

7 Morin, R.D. et al. (2008) Application of massively parallel sequencing to microRNA profiling and discovery in human embryonic stem cells. Genome Res. 18, 610–621

8 Neilsen, C.T. et al. (2012) IsomiRs--the overlooked repertoire in the dynamic microRNAome. Trends Genet. 28, 544–549

9 Ha, M. and Kim, V.N. (2014) Regulation of microRNA biogenesis. Nat. Rev. Mol. Cell Biol. 15, 509–524

10 Bofill-De Ros, X. et al. (2019) Structural Differences between Pri-miRNA Paralogs Promote Alternative Drosha Cleavage and Expand Target Repertoires. Cell Rep. 26, 447–459.e4

11 Gu, S. et al. (2012) The loop position of shRNAs and pre-miRNAs is critical for the accuracy of dicer processing in vivo. Cell 151, 900–911

12 Ma, H. et al. (2013) Lower and upper stem-single-stranded RNA junctions together determine the Drosha cleavage site. Proc Natl Acad Sci USA 110, 20687–20692

13 Bofill-De Ros, X. et al. (2019) IsomiRs: Expanding the miRNA repression toolbox beyond the seed. Biochim. Biophys. Acta Gene Regul. Mech. DOI: 10.1016/j.bbagrm.2019.03.005

14 Sheu-Gruttadauria, J. et al. (2019) Structural Basis for Target-Directed MicroRNA Degradation. Mol. Cell 75, 1243–1255.e7

15 Yang, A. et al. (2020) AGO-bound mature miRNAs are oligouridylated by TUTs and subsequently degraded by DIS3L2. Nat. Commun. 11, 2765

16 Fernandez-Valverde, S.L. et al. (2010) Dynamic isomiR regulation in Drosophila development. RNA 16, 1881–1888

17 Rüegger, S. and Großhans, H. (2012) MicroRNA turnover: when, how, and why. Trends Biochem. Sci. 37, 436–446

18 Yang, A. et al. (2019) 3’ Uridylation Confers miRNAs with Non-canonical Target Repertoires. Mol. Cell 75, 511–522.e4

19 Esquela-Kerscher, A. and Slack, F.J. (2006) Oncomirs - microRNAs with a role in cancer. Nat. Rev. Cancer 6, 259–269

20 Chu, A. et al. (2016) Large-scale profiling of microRNAs for The Cancer Genome Atlas. Nucleic Acids Res. 44, e3

21 Cancer Genome Atlas Research Network et al. (2013) The Cancer Genome Atlas Pan-Cancer analysis project. Nat. Genet. 45, 1113–1120

22 Telonis, A.G. et al. (2015) Beyond the one-locus-one-miRNA paradigm: microRNA isoforms enable deeper insights into breast cancer heterogeneity. Nucleic Acids Res. 43, 9158–9175

23 McCall, M.N. et al. (2017) Toward the human cellular microRNAome. Genome Res. 27, 1769–1781

24 Lan, C. et al. (2018) An isomiR expression panel based novel breast cancer classification approach using improved mutual information. BMC Med. Genomics 11, 118

25 Parafioriti, A. et al. (2020) Expression profiling of microRNAs and isomiRs in conventional central chondrosarcoma. Cell Death Discov. 6, 46

26 Koppers-Lalic, D. et al. (2016) Non‑invasive prostate cancer detection by measuring miRNA variants (isomiRs) in urine extracellular vesicles. Oncotarget 7, 22566–22578

27 Wang, S. et al. (2019) Tumor classification and biomarker discovery based on the 5’isomiR expression level. BMC Cancer 19, 127

28 Kozomara, A. et al. (2019) miRBase: from microRNA sequences to function. Nucleic Acids Res. 47, D155–D162

29 Fromm, B. et al. (2020) MirGeneDB 2.0: the metazoan microRNA complement. Nucleic Acids Res. 48, D132–D141

30 Zhang, Y. et al. (2016) IsomiR Bank: a research resource for tracking IsomiRs. Bioinformatics 32, 2069–2071

31 Cheng, W.-C. et al. (2013) YM500: a small RNA sequencing (smRNA-seq) database for microRNA research. Nucleic Acids Res. 41, D285–94

32 Ziemann, M. et al. (2016) Evaluation of microRNA alignment techniques. RNA 22, 1120–1138

33 Guo, L. et al. (2016) miR-isomiRExp: a web-server for the analysis of expression of miRNA at the miRNA/isomiR levels. Sci. Rep. 6, 23700

34 Bofill-De Ros, X. et al. (2019) QuagmiR: a cloud-based application for isomiR big data analytics. Bioinformatics 35, 1576–1578

35 Kim, H. et al. (2020) A Mechanism for microRNA Arm Switching Regulated by Uridylation. Mol. Cell 78, 1224–1236.e5

36 Wang, Y. and Liang, H. (2018) When micrornas meet RNA editing in cancer: A nucleotide change can make a difference. Bioessays 40,

37 Paul, D. et al. (2017) A-to-I editing in human miRNAs is enriched in seed sequence, influenced by sequence contexts and significantly hypoedited in glioblastoma multiforme. Sci. Rep. 7, 2466

38 Roberts, J.T. et al. (2018) ADAR mediated RNA editing modulates microrna targeting in human breast cancer. Processes (Basel) 6,

39 Schwartz, S. and Motorin, Y. (2017) Next-generation sequencing technologies for detection of modified nucleotides in RNAs. RNA Biol. 14, 1124–1137

40 Medina, P.P. et al. (2010) OncomiR addiction in an in vivo model of microRNA-21-induced pre-B-cell lymphoma. Nature 467, 86–90

41 Griffiths-Jones, S. et al. (2011) MicroRNA evolution by arm switching. EMBO Rep. 12, 172–177

42 Chen, L. et al. (2018) miRNA arm switching identifies novel tumour biomarkers. EBioMedicine 38, 37–46

43 Zhang, Z. et al. (2019) microRNA arm-imbalance in part from complementary targets mediated decay promotes gastric cancer progression. Nat. Commun. 10, 4397

44 Kern, F. et al. (2020) miRSwitch: detecting microRNA arm shift and switch events. Nucleic Acids Res. 48, W268–W274

45 Gu, S. et al. (2011) Thermodynamic stability of small hairpin RNAs highly influences the loading process of different mammalian Argonautes. Proc Natl Acad Sci USA 108, 9208–9213

46 Boele, J. et al. (2014) PAPD5-mediated 3’ adenylation and subsequent degradation of miR-21 is disrupted in proliferative disease. Proc Natl Acad Sci USA 111, 11467–11472

47 Shukla, S. et al. (2019) The RNase PARN Controls the Levels of Specific miRNAs that Contribute to p53 Regulation. Mol. Cell 73, 1204–1216.e4

48 Park, S.-M. et al. (2008) The miR-200 family determines the epithelial phenotype of cancer cells by targeting the E-cadherin repressors ZEB1 and ZEB2. Genes Dev. 22, 894–907

49 Wang, Y. et al. (2017) Systematic characterization of A-to-I RNA editing hotspots in microRNAs across human cancers. Genome Res. 27, 1112–1125

50 Ramírez-Moya, J. et al. (2020) ADAR1-mediated RNA editing is a novel oncogenic process in thyroid cancer and regulates miR-200 activity. Oncogene 39, 3738–3753

51 Londin, E. et al. (2015) Analysis of 13 cell types reveals evidence for the expression of numerous novel primate- and tissue-specific microRNAs. Proc Natl Acad Sci USA 112, E1106–15

